# Expression landscape of metabolic engineering enzymes in cyanobacteria

**DOI:** 10.64898/2026.04.08.717203

**Authors:** Hitesh Medipally, Anna Karlsson, Aman Dheer, Elton P. Hudson, Elias Englund

## Abstract

Photosynthetic cyanobacteria are promising platforms for sustainable chemical production, as they can convert light and CO₂ into valuable compounds. Achieving this often requires engineering cyanobacteria with non-native enzymes with strong promoters to maximize enzyme accumulation. However, despite extensive engineering efforts, the extent to which these enzymes misfold and undergo degradation in cyanobacteria remains unknown. Here, we systematically investigate the fate of recombinant proteins in *Synechocystis* sp. PCC 6803 by estimating protein loss due to protease degradation. To do this, we developed a quantitative approach that combines split-GFP reporting with inducible CRISPRi knockdown of Clp protease system, enabling estimation of portion of proteins that would otherwise be degraded. Applying this method to 103 heterologous proteins previously used in cyanobacterial metabolic engineering studies, we find that, on average, one-third of recombinant protein accumulation is lost to degradation, with some enzymes exhibiting more than 95% protein loss. Furthermore, we compare expression from identical expression constructs in *E. coli* and *Synechocystis* and find broad similarities in their protein accumulation patterns. Together, these findings provide the first quantitative overview of heterologous protein expression in cyanobacteria and identify enzymes that are suboptimal for their respective pathways, information usable to increase production titers in photosynthetic cell factories.

## Introduction

As the world moves away from fossil fuels and toward renewable production, engineering bacteria to produce fuels and chemicals offers significant potential for sustainable manufacturing. Photosynthetic cyanobacteria are especially interesting for this purpose because they use light and CO₂ as energy and carbon sources, minimizing the need to supply growth substrates, which can account for ∼70% of the total operating cost of an industrial fermenter^1^. Cyanobacteria have been engineered to produce a wide range of molecules, including fuels, plastics, and pharmaceuticals^2^. However, the technology still struggles to become economically viable, largely because product titers remain too low^3^. Achieving high titers requires high enzyme levels to redirect metabolism toward the desired product. High enzyme levels, in turn, require strong protein expression. In fact, metabolic engineering projects often rely on the strongest available promoters to pack cells with as much recombinant protein as possible.

Furthermore, studies on cyanobacteria have found a linear correlation between enzyme levels and product titers when the enzyme is the bottleneck of a pathway^4^. Overall, the viability of this technology depends on achieving high titers, and therefore requires a strong understanding of protein expression and precise control over it.

Studies on other bacteria have shown that a large percentage of heterologous proteins cannot be expressed in a soluble form. In one large-scale expression study in *E. coli*, 9,644 nonnative proteins, mostly of bacterial origin, were tested for soluble expression^5^. Among these, 25% showed no expression at all, and of the proteins that could be expressed, 30% were completely insoluble, with no detectable soluble fraction.

Similarly, another study expressed 10,167 eukaryotic proteins from *C. elegans* in *E. coli* and found that only 15% were soluble^6^. These findings highlight the challenge of expressing nonnative proteins in a host, where differences in the cellular environment, folding machinery, and molecular crowding compared to their native organism often cause misfolding and protein aggregation. The Gibbs free energy difference between the folded and unfolded states of natural proteins is only about −3 to −10 kcal/mol, roughly the same energy as a single hydrogen bond^7^. Evolution has selected for proteins to be only marginally stable because their catalytic activity relies on mechanical flexibility and overly rigid structures often reduce activity. Furthermore, overexpression of proteins negatively impacts their ability to form a soluble state. It is believed that proteins are expressed in their native host close to their solubility limit^8^. If a protein accumulates above their soluble limit, the most favorable state shifts from its soluble native form to an aggregated state. Failure to reach its native state can expose normally buried hydrophobic residues, leading to aggregation with other proteins and triggering the cellular quality control machinery to degrade the misfolded protein^9^. Therefore, small perturbation of the cellular context can change a normally folded protein to an insoluble expressed.

Although cyanobacteria, such as the unicellular model cyanobacteria *Synechocystis* sp. PCC 6803 (*Synechocystis*), have been engineered to express hundreds of proteins, the extent to which these proteins misfold and are degraded is largely unknown and rarely tested. In a study by Zhang et al. (2021), heterologous proteins were overexpressed in *Synechocystis* at high levels by fusing them to a native protein^10^.

However, as soon as the fusion was cleaved, the heterologous proteins became undetectable in the cells, due to rapid protease degradation. This rapid degradation of unstable proteins makes it difficult to determine how much of a given protein fails to fold correctly. The main protease system in cyanobacteria responsible for this degradation is the Clp protease system^11^. The Clp system is a multi-subunit AAA+ protease complex composed of an ATP-dependent chaperone component associated with a proteolytic core, where the chaperones recognize and unfold target proteins before translocating them into the degradation chamber for proteolysis. It is the primary cytoplasmic quality control protease, and several of its core components are essential, as attempts to knock them out in other cyanobacteria have failed, highlighting their critical role in maintaining protein homeostasis^12^. In addition to the Clp system, *Synechocystis* also contains the HtrA/Deg system, which targets periplasmic proteins, and FtsH, a membrane-integrated protease that acts on membrane proteins such as photosystem components^13,14^.

Our goal with this study was to systematically assess heterologous protein stability in *Synechocystis* and determine the extent to which misfolding and degradation limit protein accumulation. We developed Clp protease knockdown strains to quantify the fraction of heterologous proteins lost to degradation and applied this method to over a hundred different proteins previously used in metabolic engineering studies. This study represents the first large, systematic assessment of heterologous protein stability in cyanobacteria, providing a comprehensive resource to guide future metabolic engineering strategies and identify potential enzyme bottlenecks in heterologous pathways.

## Results

### Testing expression of stability reference proteins in *Synechocystis*

To develop a method for assessing recombinant protein expression in cyanobacteria, we selected three stability reference proteins: acylphosphatase (AcP), immunity protein 7 (Im7), and triosephosphate isomerase (Tim). These proteins were chosen because they are widely used in protein stability studies and have well characterized point mutations that affect their stability^15–17^. For each protein, we selected five single point mutations with known effect on protein stability, either stabilizing or destabilizing the structure, as measured by changes to Gibbs free energy (ΔΔG°) or protein melting temperature. Proteins with low stability have a higher chance to misfold and degrade than proteins with high stability. Therefore, our choice of using proteins with a range of different stability was a way to compare the same proteins when it has high misfolding and low misfolding. Additionally, using single amino acid variants allowed us to systematically test how protein stability influences misfolding, while keeping the coding sequences nearly identical. This allowed us to focus on factors that are intrinsic to the protein itself, and minimize variation from codon usage, mRNA stability, and other sequence-dependent effects on expression.

To be able to quantify how much protein our differently stable variants would produce, we fused the wild-type sequence and five mutant variants of AcP, Im7 and Tim to split-GFP and cloned them into a broad-host-range plasmid (**Figure 1a, b**). The split-GFP tag allows monitoring of protein expression with minimal perturbation from the tag to the native structure^18^. The resulting 18 constructs were introduced into *Synechocystis* via conjugation and protein levels were assessed by GFP fluorescence. Across all three proteins, variants with higher intrinsic stability accumulated to higher levels than less stable variants, with expression differences ranging from 2.6- to 31.5-fold between the highest and lowest expressing variants (**Figure 1c**). This trend is consistent with low stability proteins being more prone to misfolding and subsequent degradation by cellular quality control systems. Western blot analysis confirmed that differences in fluorescence corresponded to differences in protein abundance, rather than inaccessibility of the GFP tag (**Figure 1d**).

**Figure 1.**
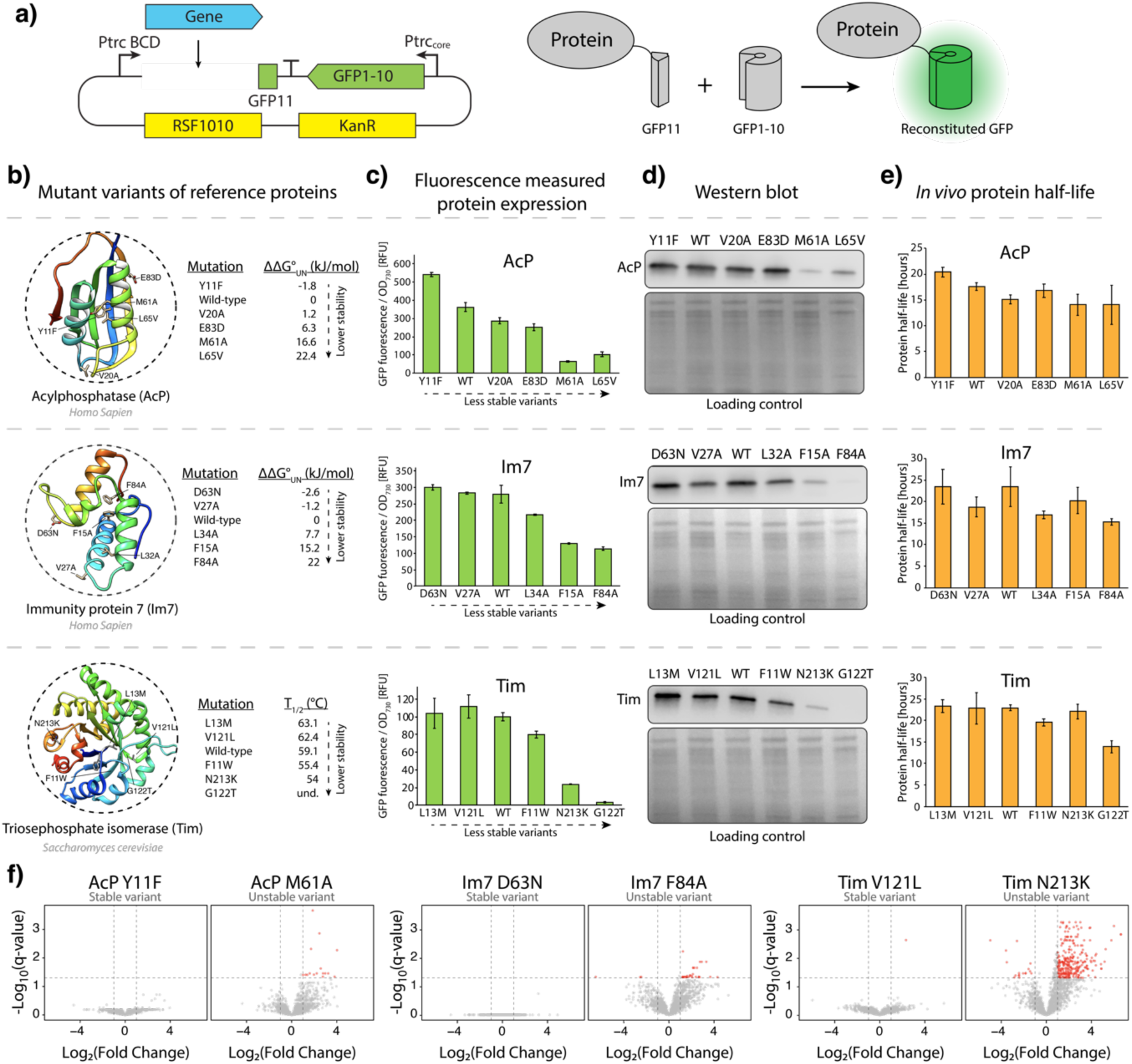
Expression of reference proteins with different stability in *Synechocystis*. a) Gene construct design where protein expression is monitored by split-GFP. One part of GFP (GFP11) is attached to the protein of interest and a nonfluorescent GFP lacking a beta strand (GFP1-10) is expressed separately^18^. Only a reconstituted GFP gives fluorescence. Reconstitution figure based on Kamiyama et al. (2016)^19^. b) Protein structure of reference proteins AcP (AlphaFold prediction), Im7 (1AYI) and Tim (1YP1). Positions of mutations are marked on their structures. Values for changes to stability and melting temperature was experimentally determined in previous studies for AcP^20^, Im7^15,16,21^ and Tim^17^. ΔΔG°_UN_ values are defined as ΔG°_UN_ (mutant) - ΔG°_UN_ (wild-type). und. = undetermined Tm, due to the variant being unpurifiable. c) GFP fluorescence measurements from *Synechocystis*. Mutant variants are ordered by stability. d) Western blot analysis against N-terminal strep-tag of total proteins, including both soluble and insoluble fractions. Coomassie stained SDS-PAGE was used as equal sample loading control. e) Antibiotic chase assay to measure protein half-life *in vivo*. Chloramphenicol (200 μg/mL) and spectinomycin (200 μg/mL) were added to growing cultures and decrease in fluorescence was used to determine protein half-life of AcP, Im7 and Tim variants. Error bars represent the standard deviation of three biological replicates. f) Volcano plots of proteomic profiles for stable and unstable variants of the reference proteins. Protein abundances were compared to an empty-vector control. Differentially abundant proteins (red dots) were defined as those with an adjusted p-value < 0.05 and a log_2_ fold change of ≥1 or ≤-1.

Next, we investigated whether variation in protein half-life could explain the differences in protein accumulation between the variants. Translation was halted using two ribosome targeting antibiotics, and the decline in fluorescence was monitored as a measure of protein degradation. While low stability variants tended to have shorter half-lives (**Figure 1e**), these differences were insufficient to account for the large variation in protein levels seen in **Figure 1c**. Because fluorescence is only emitted by fully synthesized and mature proteins, this assay measures the degradation of the mature protein pool and does not capture misfolding or degradation events that occur during protein synthesis. The discrepancy between protein half-life and steady-state accumulation therefore suggests that a substantial fraction of protein quality control occurs co-translationally, with unstable variants being targeted for degradation before reaching their fully folded, fluorescent state.

To investigate how expression of proteins with different intrinsic stabilities affected the host cell, we performed quantitative proteomic analysis on *Synechocystis* strains expressing high- and low-stability variants of AcP, Im7, and Tim. For each variant, protein abundance was compared to an empty vector control strain to identify changes in the cellular proteome. Proteins with an adjusted p-value < 0.05 (corrected for multiple hypothesis testing) and log_2_ fold change ≥1 were considered significantly altered. Expression of the high-stability variants resulted in minimal changes, with no significantly altered proteins identified for two of the three variants and only a single significantly changed protein for the third (**Figure 1f**, Supplementary File 1). In contrast, all three low-stability variants induced some degree of changes in protein abundance, with the Tim N_213K_ variant showing the largest response, affecting nearly 300 proteins.

These changes included proteins involved in translation and amino acid biosynthesis, components of photosynthetic complexes, and enzymes linked to glycogen metabolism. In addition, several proteins associated with protein quality control were among the upregulated set, including components of the Clp protease system. These findings show that expression of unstable variants, in contrast to stable variants, triggers a broad cellular response.

### Creating protease knockdown strains

A challenge for developing methods to assess soluble expression is that rapidly degraded proteins are difficult to quantify. Inspired by the *E. coli* BL21 (DE3) system, where the main quality control protease Lon is knocked out to allow accumulation of otherwise degraded proteins, we hypothesized that targeted knockdowns of native proteases in *Synechocystis* could similarly prevent degradation of misfolded proteins. By reducing protease activity, the additional proteins that accumulated could serve as a way for estimating degradation and solubility of heterologous proteins.

*Synechocystis* does not have Lon proteases. Instead, they use Clp proteases to process cytosolic proteins (**Figure 2a**)^11^. They also have Deg/HtrA system associated with periplasmic proteins, and FtsH proteases that are associated with thylakoid membrane protein turnover^13,14^. Because several parts of the Clp systems are essential for cell survival, creating complete knockouts of all components is not viable^12^. Instead, we used a previously developed CRISPRi approach to create inducible knockdowns^22^. The CRISPRi method uses dCas9 integrated into the genome in the *psbA1* gene, acting as a neutral site, together with two guide RNAs (gRNA) under the control of a TetR regulated promoter (**Figure 2b**). Gene knockdown is thus induced by addition of anhydrotetracycline (aTc), which activates gRNA and dCas9 expression. As two gRNA are used, each strain could be designed to simultaneously target two genes for knockdown.

**Figure 2.**
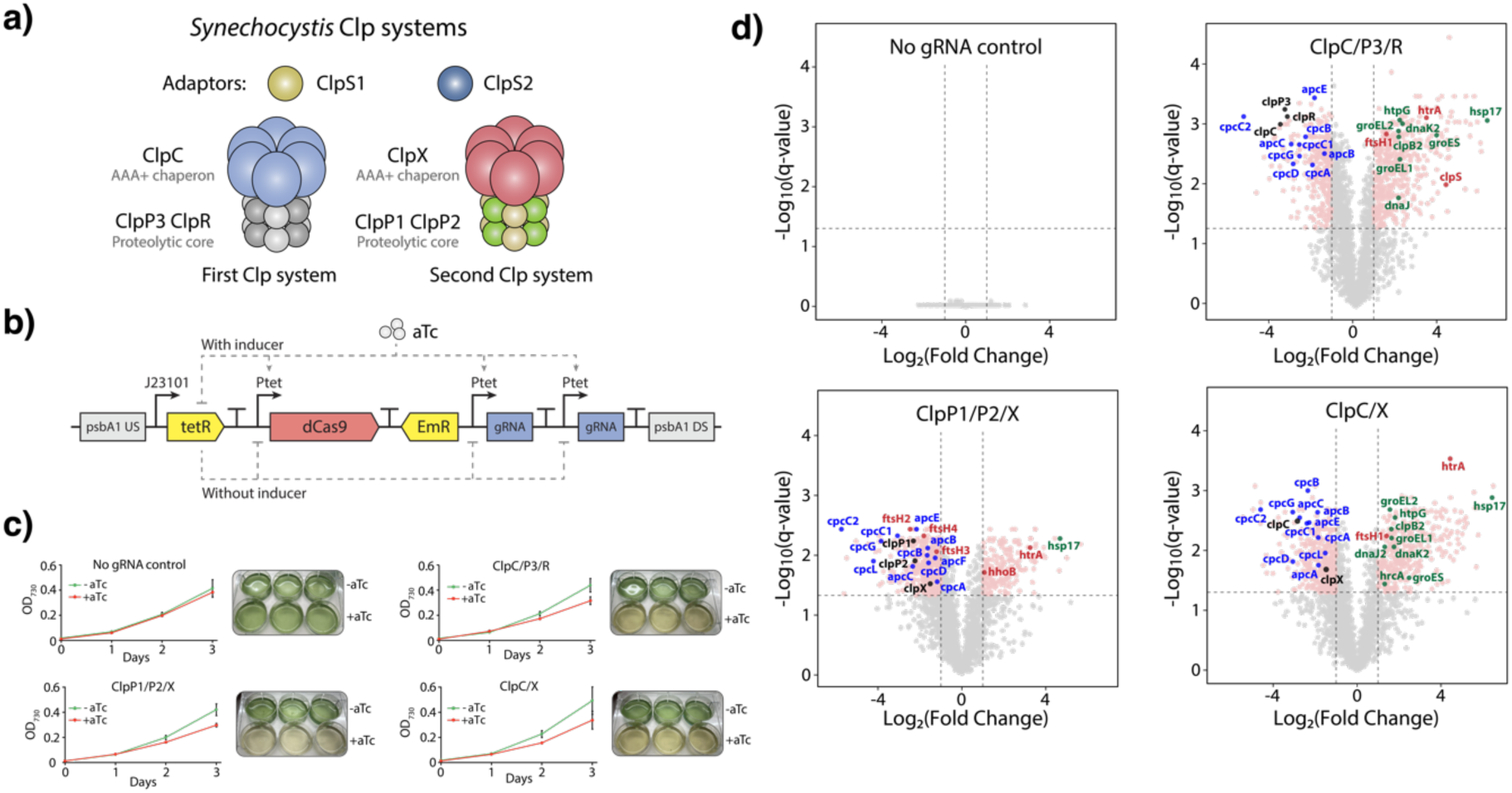
CRISPRi knockdown of proteases in *Synechocystis*. a) Clp protease systems of *Synechocystis*. We call ClpP3/R/C as the first system and ClpP1/P2/X as the second system to simplify referring to them in this manuscript. Figure inspired by Bouchnak C Wijk (2020)^11^. b) Genetic construction for CRISPRi knock-down. dCas9 and two guide RNAs (gRNAs) targeting two different proteases are under the control of aTc inducible promoters positioned in the *psbA1* gene that acts as a neutral site.) Growth curve of Clp CRISPRi strains, growing with and without knockdown inducer aTc (1.25 μg/mL). Inserted pictures show chlorotic phenotype at day 3. Optical density was measured using a plate reader which gives OD_730_ values ∼5 times lower than when measured with a standard 1 cm pathlength cuvette spectrophotometer. Error bars represent standard deviation of three biological replicates. d) Volcano plot from proteomic analysis of Clp knockdown strains, comparing protein abundance in the same strains with and without aTc induced knockdown, two days after induction. Targets of knockdown are color coded black, phycobiliproteins are blue, proteases are red and chaperones/heat shock proteins are green. Results are based on three biological replicates. Growth curves and volcano plots from ClpS1/S2 and Deg/HtrA knockdown strains are available in Supplementary Figures 1 and 2.

We selected components of the Clp and Deg/HtrA protease systems as targets for knockdown, including the proteolytic core, chaperones, and adaptors of the Clp system (**Table 1**). In total, seven protease knockdown strains were generated, including two targeting the Deg/HtrA system, with two genes targeted per strain. In some strains, downstream genes in operons were also silenced, as for the ClpC/P3/R strain, where ClpP3 was the primary target for one of the gRNAs and ClpR was silenced because of its downstream position. All strains could be generated except one that targeted the proteolytic core of both Clp systems (ClpP1 and ClpP3/R), which could not be obtained despite repeated attempts. This highlights the importance of Clp, as even the uninduced expression of the dCas9 knockdown system using a weak and tightly regulated promoter (P_L22_ variant^23^) was sufficient to be lethal to the cells.

**Table 1.** Result of proteomic analysis of *Synechocystis* protease knockdown strains. Strains growing with and without CRISPRi inducer aTc were compared to identify proteins with significant changes to their abundance (log_2_ fold change ≥1, adjusted p-value < 0.05). Knockdown of target genes were significant if p-value < 0.05 (not adjusted p-value). CRISPRi repression fold changed is in base 10.

| Strain/gRNA targets | Intended target | Proteins detected | Differently expressed proteins | CRISPRi repression (fold change) |
| --- | --- | --- | --- | --- |
| No gRNA control | None | 2384 | 0 | N/A |
| ClpC + ClpP3/R | First Clp system | 2521 | 797 | ClpC: -12.3<br>ClpR: -9.8<br>ClpP3: -10.6 |
| ClpP1 + ClpP2/X | Second Clp system | 2366 | 626 | ClpP1: -4.7<br>ClpP2: -4.7<br>ClpX: -2.8 |
| ClpC + ClpX | Chaperones of both Clp systems | 2465 | 749 | ClpC: -7.2<br>ClpX: -2.8 |
| ClpP1 + ClpP3/R | Proteolytic core of both Clp systems | N/A | N/A | Unable to generate strain |
| ClpS1 + ClpS2 | Clp adaptors | 2410 | 0 | ClpS1: Undetected<br>ClpS2: -4.3 |
| HhoA + HtrA | Deg/HtrA system | 2372 | 1 | HhoA: -12.9<br>HtrA: Nonsignificant |
| HhoB + HtrA | Deg/HtrA system | 2386 | 4 | HhoB: -6.8<br>HtrA: -1.9 |

The six successfully generated knockdown strains, along with a no gRNA control, were tested for growth with and without inducer. Strains targeting ClpC + ClpP3/R (core and chaperone of one Clp system), ClpP1 + ClpP2/X (core and chaperone of the other Clp system), and ClpC + ClpX (chaperones of both systems) showed growth reduction and a bleaching phenotype upon inducing dCas9 knockdown (**Figure 3c**). Interestingly, when ClpP1, ClpP2 and ClpX were knocked out in *Synechococcus* sp. PCC 7942, none of those strains was reported having bleaching^12^. Our knockdown of Clp adaptors ClpS1 + ClpS2, as well as the two Deg/HtrA strains, showed no growth defects (Supplementary Figure 1). This is consistent with previous reports on Deg/HtrA knockouts in *Synechocystis*, which also showed no effect on growth^24^.

**Figure 3.**
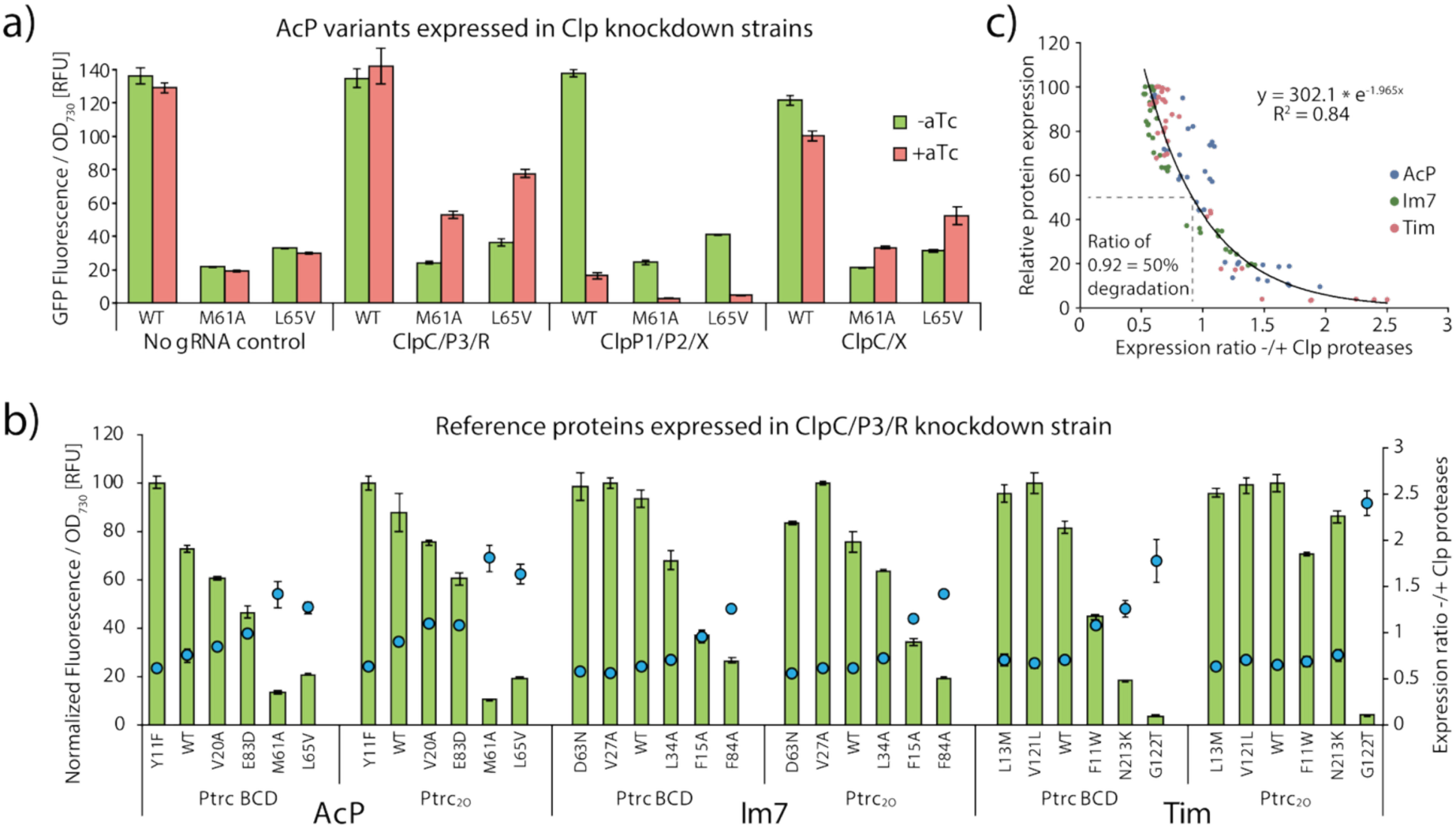
Expression of reference proteins with varying stability in *Synechocystis* protease knockdown strains. a) Expression of AcP variants in protease knockdown strains. CRISPRi knockdown was induced by aTc. b) Expression of AcP, Im7 and Tim variants using Ptrc BCD or Ptrc_20_ as promoters in ClpC/P3/R strain. The left y-axis shows fluorescence (green bars) normalized to the highest expressing variant within each data set, and right y-axis represents expression ratio (blue dots) calculated as fluorescence with aTc induction and Clp knockdown divided by fluorescence without induction and with unaltered Clp levels (−/+ Clp, corresponding to the red/green bar ratio in Fig. 3a). Error bars represent standard deviation of three biological replicates. c) Relationship between expression ratio −/+ Clp proteases in ClpC/P3/R strain and relative protein expression for individual measurements from Fig 3b. The data were used to establish a correlation factor allowing conversion of −/+ Clp ratio values into estimations of protein loss due to degradation.

### Proteomic analysis of protease knockdown strains

To understand the cellular response to the protease knockdowns, we performed proteomic analysis on all strains, comparing changes to proteome of cultures growing with inducer (aTc) to cultures growing without (**Figure 2d**, Supplementary Files 2-9).

Across all strains, our method detected an average of 2410 proteins. We saw a significant reduction in protein levels of the CRISPRi targeted proteases, ranging from 1.9-fold to 12.3-fold reduction, except for ClpS1, which was undetected, and HhoA which showed no significant reduction in one of the strains (**Table 1**). A substantial proportion of detected proteins showed altered abundance in the ClpC/P3/R, ClpP1/P2/R, and ClpC/X strains, corresponding to 32%, 26%, and 30% of detected proteins, respectively. In contrast, the negative control and ClpS1/S2 had no significant changes, while Deg/HtrA knockdown only affected a handful of proteins (Supplementary Figure 2).

The most pronounced effects were observed in the strains targeting proteolytic cores or chaperones of the Clp systems. These strains exhibited upregulation of many chaperones, with heat shock protein Hsp17 showing the largest increase of all proteins, accompanied by elevated levels of stress sigma factors SigB and SigE. Several proteases were also upregulated, likely as a compensatory response to reduced Clp activity or an increase in misfolded proteins. Phycobiliproteins were consistently downregulated, reflecting the observed bleaching phenotype. We verified these results by extracting and measuring pigment spectroscopically, and found that together with phycocyanin pigments, chlorophyll content was also reduced, while carotenoids were unaffected in all strains except ClpP1/2/X (Supplementary Figure 3).

Other than pigments, proteases and chaperones, there was no clear, easily explained consistent pattern of up- or downregulation of cellular systems for the three Clp knockdown strains. Nitrogen starvation produces a similar bleaching phenotype to that observed here. That response is mediated by NblA1, NblA2 and NblD^25,26^, but we could not detect these proteins with our method. Comparing our results with a previous RNA-seq study on nitrogen starvation in *Synechocystis*^25^, we found similarities, like upregulation of nitrate transporters NrtA-D, but also many differences, like downregulation of master regulators PII (GlnB) and oxidative pentose phosphate proteins Zwf and Gnd, indicating that this response is different from the nitrogen starvation response.

Interestingly, proteins previously identified as ClpX interaction partners in a *Synechocystis* pull-down study showed a predominant decrease in abundance following ClpX knockdown^27^. If ClpX primarily binds proteins targeted for degradation, reduced ClpX activity would instead be expected to result in the accumulation of these proteins. However, the opposite trend was observed, with the strongest effect in the ClpP1/P2/X strain, where 129 known ClpX interactors decreased in abundance compared with only 9 that increased (Supplementary Table 1). This contrasts with the overall proteome-wide response for that strain, where the majority of significantly altered proteins showed increased abundance (436 increased vs. 190 decreased).

Many proteins from carbon fixation and the photosystems were also affected by the Clp knockdown. The large subunit of RuBisCO, FNR and several components of PSI were all downregulated. However, the strains had reduced growth upon Clp knockdown.

Therefore, these effects could be due to a lower need for fixed carbon and light energy. Overall, it is difficult to know if the proteomic changes observed across the Clp knockdown strains are due to targets of Clp escaping degradation, due to the accumulation of misfolded proteins, or due to the slower growth of the strains.

### Expression patterns of reference proteins in protease knockdown strains

We next tested whether the protease knockdown strains would accumulate unstable proteins, allowing us to measure what would otherwise be degraded. We hypothesized that low stability protein variants would show the largest increase in abundance upon protease knockdown, whereas high stability variants would remain largely unaffected. To test this, we introduced three AcP variants, one high stability (AcP_WT_) and two low stability (AcP_M61A_ and AcP_L65W_) variants, into all seven CRISPRi strains, generating 21 new strains. All strains were grown with or without aTc induction, and the difference in protein accumulation was measured using the split-GFP tag.

In the ClpC/P3/R knockdown strain, high stability AcP_wt_ showed no change in protein levels from the knockdown while the low stability AcP_M61A_ and AcP_L65W_ more than doubled in protein accumulation relative to the uninduced controls, in accordance with our hypothesis (**Figure 3a**). Results from the ClpC/X strain produced a similar pattern, while ClpP1/P2/X had lower expression for all variants, possibly due to loss of cell viability. No significant change in protein expression was observed upon ClpS1/S2 and Deg/HtrA knockdown (Supplementary Figure 4).

Based on these results, we focused on the ClpC/P3/R strain, which showed the most promising response. We introduced the full set of reference proteins and constructed 18 new expression plasmids, in which the Ptrc BCD promoter was changed to Ptrc_2O_. Although both promoters are derived from Ptrc, they have been reported to yield substantially different expression levels^28,29^, allowing us to evaluate whether promoter choice influences the behavior of the reference protein variants.

Using both the new Ptrc_2O_ constructs and the previously generated Ptrc BCD constructs, we measured protein expression in the presence and absence of Clp knockdown and calculated a −/+ Clp ratio for each variant. The ratio was defined as protein accumulation in the knockdown condition (−Clp, +aTc) divided by accumulation in the control condition (+Clp, no induction). A ratio above 1 therefore indicates that protein levels increase when Clp activity is reduced. Across AcP, Im7, and Tim, a consistent relationship was observed between variant stability and their −/+ Clp ratio (**Figure 3b**).

Variants with lower expression levels exhibited higher −/+ Clp ratios, suggesting that the protein is normally degraded by Clp. Overall, less stable variants showed ratios above 1, while highly stable variants had ratios around 0.5-0.7. For example, highly stable variant AcP_Y11F_ had a ratio of 0.6, whereas the less stable AcP_M61A_ had a ratio of 1.4, while only accumulating 13% of the protein that AcP_Y11F_ expressed. The low ratio value of stable variants could be due to the Clp knockdown causing cellular stress, which was evident by slower growth and pigment bleaching.

To be able to use the −/+ Clp ratio to estimate the extent of protein degradation, we used the reference protein dataset to establish a relationship between ratio values and protein loss. For each reference protein, the highest expressing variant was assumed to be minimally affected by Clp-mediated degradation and was used as a zero-degradation reference. Lower expression levels in the remaining variants were then interpreted as protein loss due to degradation and misfolding. For example, AcP_M61A_ accumulated to only 13% of the level of AcP_Y11F_, corresponding to an estimated 87% protein loss. Using this approach, expression normalized to the highest-expressing variant was plotted against the corresponding −/+ Clp ratios, yielding a fitted curve (**Figure 3c**, R^2^ = 0.84). Based on this relationship, a −/+ Clp ratio of 0.92 corresponds to approximately 50% protein loss, whereas a ratio of 1.74 corresponds to 90% protein loss.

### Estimating degradation of mevalonate pathway enzymes

To test our method for assessing degradation during recombinant protein expression, we first applied it to a small dataset. The goal was to simulate a metabolic engineering project in which genes previously optimized for expression in *E. coli* are transferred to cyanobacteria. We selected the mevalonate pathway, a non-native pathway for most bacteria, including *Synechocystis*, comprising of seven genes in total. The plasmid pMBIS^30^, commonly used to enhance terpenoid production in *E. coli*, was obtained from Addgene, and the pathway genes were cloned into our Ptrc_2O_ driven expression vector before introduction into the ClpC/P3/R strain. Expression analysis revealed that one enzyme, MK, exhibited very low expression and a −/+ Clp ratio of 1.8, indicating that more than 90% of the enzyme was degraded (**Figure 4a**). The remaining six enzymes showed an average degradation ratio corresponding to approximately 27% protein loss, while exhibiting substantially higher expression levels than MK. These results indicate that even highly accumulated enzymes remain subject to some degradation by cellular protein quality control mechanisms.

**Figure 4.**
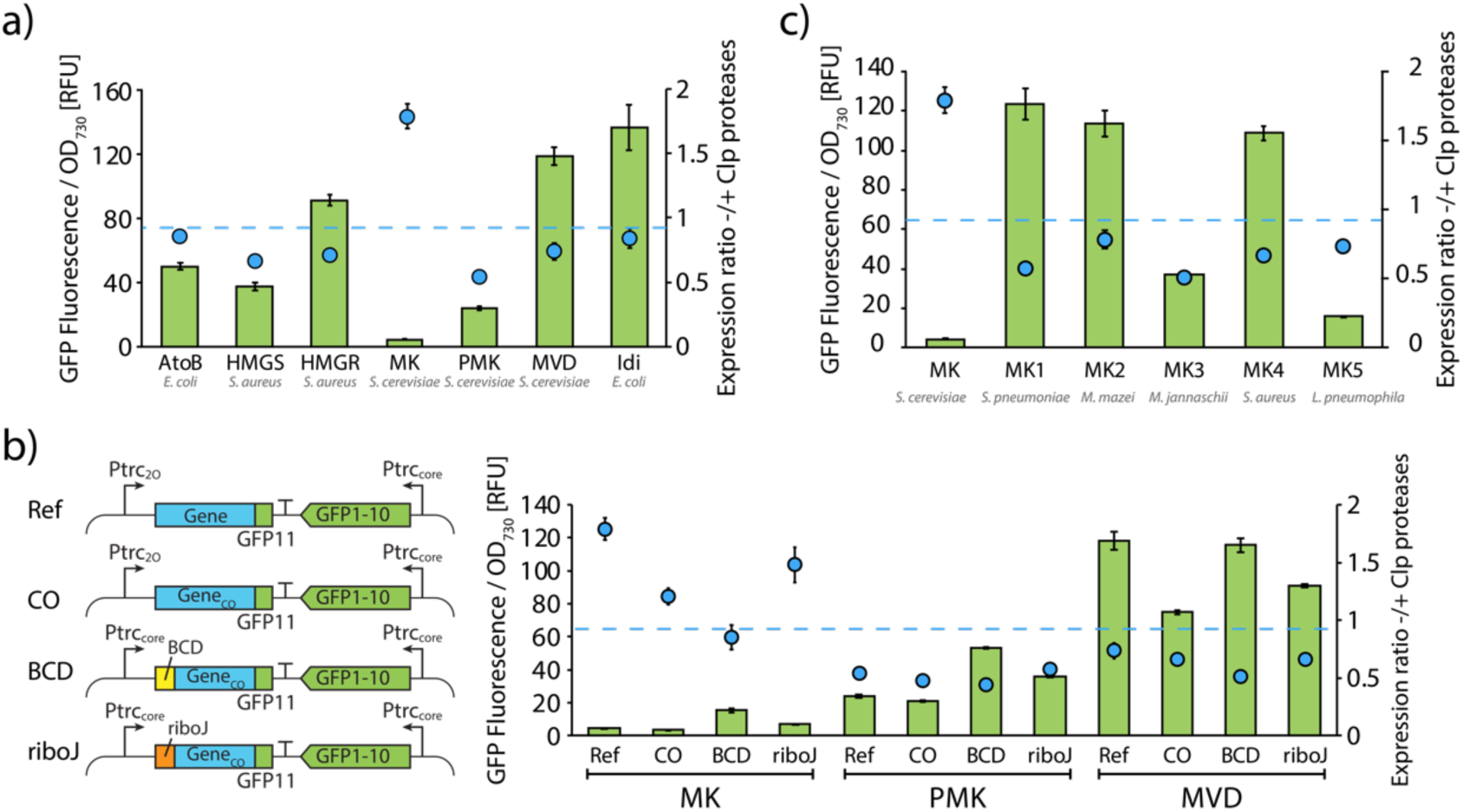
Expression of mevalonate pathway enzymes and estimated protein loss, using ClpC/P3/R knockdown strain in *Synechocystis*. a) Expression of mevalonate pathway enzymes. b) Expression and degradation of three mevalonate pathway enzymes using four genetic expression designs. Left side insert show differences between Ref, CO, BCD and riboJ genetic constructs. Ref = MK, PMK or MVD construct from Fig. 4a, BCD = bicistronic design, CO = codon optimized. c) Testing expression and degradation of MK homologs from different organisms. Green bars represent fluorescence (left axis), blue dots display the difference in fluorescence with and without Clp knockdown induced (right axis). Dashed blue line shows −/+ Clp ratio corresponding to 50% of protein loss. Error bars represent standard deviation of three biological replicates.

Next, we examined if expression levels could be improved, and degradation reduced through changes to the genetic constructs. We selected MK, PMK and MVD as they had low, medium and high expression in our measurements, and those genes were codon optimized from *E. coli* to *Synechocystis* codon usage. In addition to the original expression plasmid using Ptrc_2O_, we created new versions with expression driven by Ptrc BCD and Ptrc riboJ. Both BCD and riboJ are genetic insulators that are meant to reduce variability in gene expression^31,32^, and they have previously been used to improve protein expression in cyanobacteria^28,29^. Incorporation of a BCD increased MK expression approximately fourfold and reduced the −/+ Clp ratio from 1.8 to 0.9, corresponding to a decrease in estimated protein loss from 91% to 43% (**Figure 4b**). Although BCD elements function by enhancing ribosome binding site availability for the ribosome, there have been reports of BCDs improving soluble protein expression^33^. Codon optimization of MK also lowered the −/+ Clp ratio, despite no detectable change in overall expression levels. Consistent with this, previous studies have found that codon optimization can help with the soluble expression of proteins^34^. In contrast to MK, alterations to PMK and MVD expression constructs produced comparatively modest effects on both expression and degradation.

While modifying the genetic elements of MK had some effect on expression, protein levels were overall still low. We therefore tested if replacing MK with homologs from other organisms would have a greater impact on protein expression than modifying expression elements. Five MK homologs from different organisms were tested and all displayed higher expression levels and lower degradation compared with the original *S. cerevisiae* MK (**Figure 4c**). These results suggest that substituting poorly expressing pathway enzymes with homologous variants may be a more effective strategy than optimizing genetic expression elements.

### Measuring enzymes from metabolic engineering studies on cyanobacteria

With a method to assess the degradation of proteins established, we applied it on a larger data set of enzymes previously used for metabolic engineering of cyanobacteria. We surveyed the metabolic engineering literature to identify enzymes from as many previously engineered pathways in cyanobacteria as possible, including pathways for alcohols, sugars, terpenoids, fatty acids, and amino acid derivatives. When multiple analogous enzymes were reported for a pathway, we chose the “best in test” enzyme that achieved the highest titers. Proteins used as genetic tools were also included, such as antibiotic resistance proteins, transcription factors, fluorescent proteins and CRISPR-Cas proteins. Membrane-associated and periplasmic proteins, native *Synechocystis* proteins, and proteins requiring additional subunits or maturation processes were excluded from consideration. In total, 107 proteins were selected for our study, 86 metabolic enzymes and 21 proteins used for genetic engineering, four of which could not be expressed due to toxicity. See Supplementary File 10 for a complete list of measurement results, genes included in the study, and their corresponding sequences.

Two things were measured in our large data set to evaluate functional expression; expression as measured by split-GFP fluorescence and degradation as measured by −/+ Clp ratio. Results are summarized in **Figure 5**. Among all metabolic enzymes tested, the average degradation rate was 34%, and 24/84 enzymes had a −/+ Clp ratio value above 0.92, suggesting that half or more of protein expression is lost to degradation. This is despite our selection of enzymes being biased, as we only included enzymes from studies that successfully demonstrated production, which depends on active enzyme expression.

**Figure 5.**
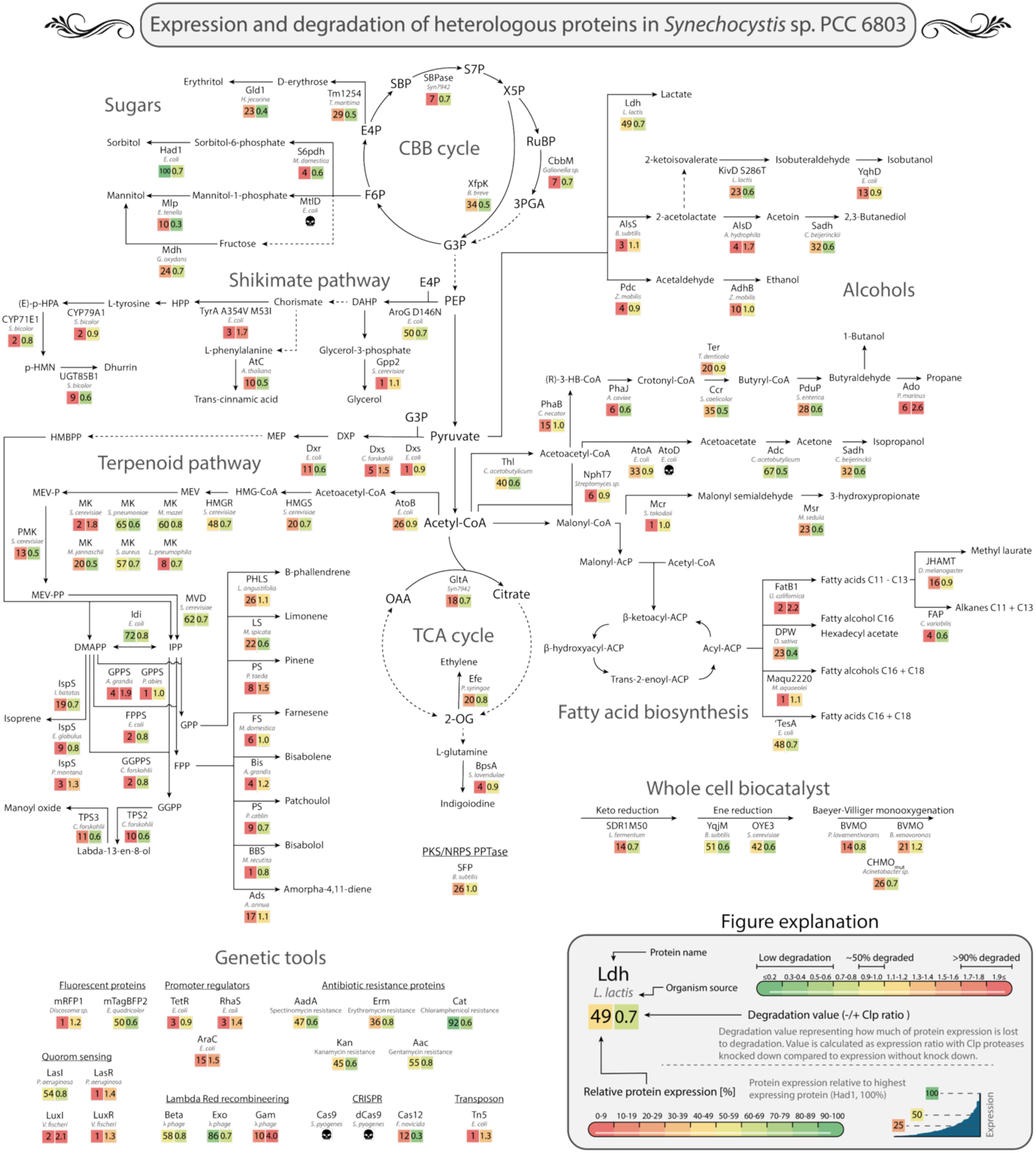
Estimating degradation of enzymes previously used for metabolic engineering of cyanobacteria. 103 heterologous proteins were measured for their expression using Ptrc_2O_, as measured by split-GFP tag, and their degradation, as measured by −/+ Clp ratio in the ClpC/P3/R strain. All expression values are normalized to the highest expressing protein, Had1. Solid lines show individual enzymatic steps, dashed lines represent multiple steps. Skull icons denote proteins that could not be measured due to presumed toxicity. Results are based on three biological replicates. A full list of results, including standard deviation, is available as Supplementary File 10. Abbreviations: SBP = sedoheptulose-1,7-bisphosphate; S7P = sedoheptulose-7-phosphate; RuBP = ribulose-1,5-bisphosphate; 3PGA = 3-phosphoglycerate; G3P = glyceraldehyde-3-phosphate; F6P = fructose-6-phosphate; E4P = erythrose-4-phosphate; (R)-3-HB-CoA = (R)-3-hydroxybutyryl-coenzyme A; PEP = phosphoenolpyruvate; 2-OG = 2-oxoglutarate; OAA = oxaloacetate; DAHP = 3-deoxy-D-arabino-heptulosonate-7-phosphate; HPP = 4-hydroxyphenylpyruvate; (E)-p-HPA = (E)-p-hydroxypenylacetaldoxime; p-HMN = p-hydroxymandelonitrile; DXP = 1-deoxy-D-xylulose-5-phosphate; MEP = 2-C-methyl-D-erythritol-4-phosphate; HMBPP = (E)-4-hydroxy-3-methyl-but-2-enyl diphosphate; HMG-CoA = 3-hydroxy-3-methylglutaryl-coenzyme A; MEV = mevalonate; MEV-P = mevalonate-5-phosphate; MEV-PP = mevalonate-5-diphosphate; DMAPP = dimethylallyl diphosphate; IPP = isopentenyl diphosphate; GPP = geranyl diphosphate; FPP = farnesyl diphosphate; GGPP = geranylgeranyl diphosphate.

Most pathway contain at least one enzyme with significant degradation. For example, the butanol pathway, which was thoroughly optimized for *Synechocystis* in Lu et al. (2019) to reach titers of 4.8 g/L^28^, used PhaB from *C. necator*, which in our assay had 56% degradation. Enzymes in the terpenoid biosynthesis pathway had many enzymes with poor expression and/or high degradation. This included all four tested GPPS, FPPS and GGPPS enzymes, as well as most of the terpene synthases. Some enzymes like AlsD from the 2,3,butanediol pathway, Ado from the propane pathway, and FatB1used for fatty acid synthesis, had −/+ Clp ratio of 1,7, 2.6 and 2.2 respectively, corresponding to more than 90% degradation. These results suggest that those pathways could be improved by selecting alternative enzymes. Of the proteins used as genetic tools, all five tested antibiotic resistance proteins were well expressed while transcription factors AraC, TetR and RhaS, all had high degradation rates. These results are consistent with studies from *E. coli* showing that transcriptional regulators have a higher turnover to allow rapid responses to changing conditions^35^.

We analyzed our entire dataset to identify factors influencing expression and degradation. We found a moderate negative correlation between protein expressed levels and their −/+ Clp ratio (Spearman ρ = −0.55, p < 10^−6^), meaning that proteins that are not degraded are more likely to accumulate to higher levels. This effect could be seen for the top 20 expressing enzymes, where none had a −/+ Clp ratio of >0.9 while the bottom 20 expressing enzymes had 15 out of 20 with >0.9, highlighting that high expressing enzymes are mainly soluble expressed while low expressing have significant degradation by Clp. We also found a small but significant correlation with the size of the protein and lower expression (ρ = −0.27, p = 0.006), while there was no such correlation for protein size and −/+ Clp ratio. Additionally, we found no evidence that the N-end rule, which associates specific N-terminal residues with higher degradation rates^11^, affected protein stability under our experimental conditions. Proteins with destabilizing N-terminal amino acids (F, L, W, Y, R, K)^36^ had no significant difference in expression or Clp ratio compared to other proteins, as determined by a two tailed unpaired t-test on log_10_ transformed data. There was also no significant difference in expression or Clp ratio based on protein origin (eukaryotic or bacterial), predicted pI, fraction of charged amino acids, or GRAVY score.

Next, we investigated whether proteins produced under Clp knockdown conditions remained enzymatically active. We measured activity from lysed cell extracts of strains expressing three Baeyer-Villiger monooxygenases which all catalyze the oxidation of the same cyclohexanone substrate. BVMO_xeno_ and BVMO_parvi_ both lost activity above background levels upon Clp knockdown, while CHMO_mut_, which had a Clp ratio indicative of low Clp-dependent degradation, maintained activity proportional to its protein abundance after Clp knockdown (**Figure 6**). These results indicate that reducing Clp-mediated degradation does not necessarily result in increased levels of active enzyme. This contrasts with observations in *E. coli*, where proteins accumulating after protease removal largely retain enzymatic activity^37^.

**Figure 6.**
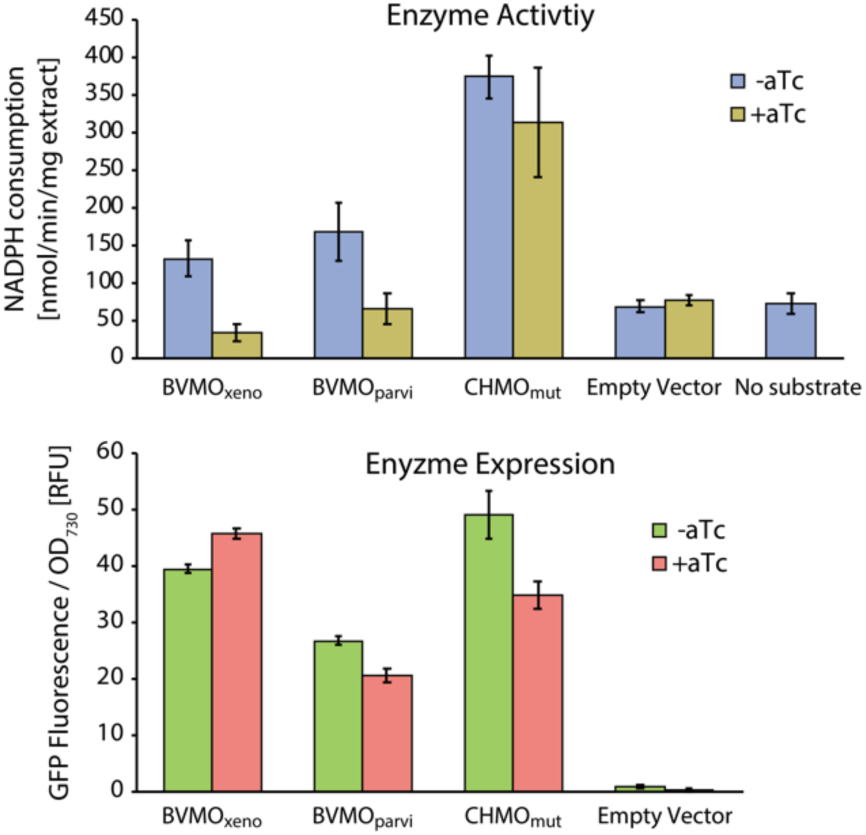
Effect of Clp KD on enzyme activity. Activity and expression of BVMO_xeno_, BVMO_parvi_ and CHMO_mut_ in ClpC/P3/R *Synechocystis* strain was measured using lysed cell extract and fluorescence. Depletion of NADPH was measured at A340 upon addition of the cyclohexanone substrate. Empty vector and No substrate (cyclohexanone) samples were controls of background NADPH depletion reactions from cell extract. Error bars represent standard deviation of three biological replicates.

Lastly, we compared protein expression between *E. coli* and *Synechocystis*. Because the expression plasmids function in both hosts, the same set of constructs tested in *Synechocystis* could also be measured in *E. coli*. Across 138 proteins, expression levels showed a significant positive correlation between the two organisms (Spearman ρ = 0.69, p < 0.0001, **Figure 7**, Supplementary File 11), indicating that proteins that accumulate well in *E. coli* generally also accumulate well in *Synechocystis*. Despite this overall correlation, several proteins deviated strongly from the general trend. For example, TetR was among the highest expressed proteins in *E. coli* but accumulated poorly in *Synechocystis*, while MK from *S. pneumoniae* and Ccr accumulated substantially better in *Synechocystis* than in *E. coli*. Together, these results suggest that while *E. coli* expression can serve as a useful predictor of protein accumulation in *Synechocystis*, individual proteins may respond differently depending on host context.

**Figure 7.**
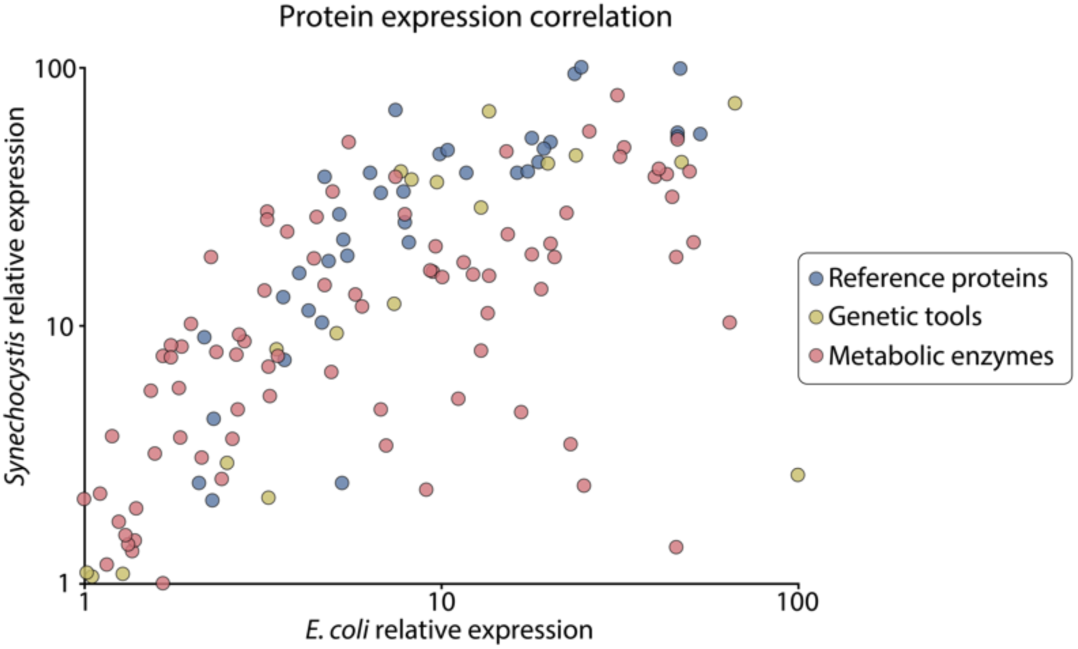
Relative protein expression in *E. coli* and *Synechocystis* of expression plasmids from Fig 3b and Fig 5. Identical expression plasmids were tested in both hosts, and protein expression was quantified as GFP fluorescence normalized to OD_600_ (*E. coli*) or OD_730_ (*Synechocystis*). Expression values were scaled relative to the lowest- and highest-expressing proteins, which were assigned values of 1 and 100, respectively.

## Discussion

Cyanobacteria have been genetically engineered with foreign genes since at least the early 1970s^38^. Despite this long history, our understanding of recombinant protein expression in cyanobacteria remains limited. Here, we systematically investigate protein expression and stability to determine the extent to which enzymes commonly selected for metabolic pathway engineering may be constrained by protein instability. This knowledge can be used to identify enzymes acting as metabolic bottlenecks and to reduce cellular resources wasted on misfolded proteins.

Testing the soluble expression of proteins in cyanobacteria presents several challenges. A major issue is that misfolded proteins are actively degraded by proteases, meaning that standard protein detection methods only capture proteins that successfully fold and accumulate. Although protein half-life can be measured, that does not capture how much protein is lost during synthesis, where most protein quality control occurs. A second challenge is distinguishing whether low protein accumulation reflects intrinsic properties of the protein or differences in transcription and translation. Because protein expression is highly sequence dependent^39^, low abundance may result from inefficient gene expression rather than protein instability, making degradation difficult to infer from protein levels alone.

To circumvent those challenges, we developed a method based on knocking down Clp proteases and measuring protein that would otherwise be degraded. Removing proteases is a well-established strategy that has been used to improve the recovery of recombinant proteins in several organisms^40,41^. It is also used when estimating soluble expression in *E. coli*, where protease-deficient strains are used to reduce degradation and allow recombinant proteins to accumulate, after which soluble and insoluble fractions are separated by centrifugation and visualized by SDS-PAGE^42^. For example, Price et al. (2011) tested the solubility of 9,644 proteins in *E. coli* by comparing total and soluble fractions on protein gels^5^. Our method, however, compares the fluorescence of GFP tagged proteins in two cultures, one with the knockdown induced and one uninduced, which is arguably a faster and less laborious process.

To isolate the effect of intrinsic protein stability on expression, we used reference proteins containing single point mutation variants. As the expression plasmids were genetically almost identical, any differences in protein accumulation were attributed to the variants’ ability to fold into soluble proteins. Testing the reference proteins in the ClpC/P3/R strain showed a negative correlation between the stability of a protein and the additional protein accumulated, which was used to create a correlation factor. To create this correlation factor, we assumed that the highest-expressing variants of AcP, Im7, and Tim experienced negligible Clp degradation, and that their expression represented 100% realized expression. This assumption might not be correct, especially for AcP, since the one mutant variant that increased stability, AcP_Y11F_, increased expression by 49%. This large increase suggests that the wild-type sequence is not well soluble in *Synechocystis,* and that additional mutations might improve solubility even further. However, this potential issue with our assumptions makes our estimation err on the conservative side, as setting AcP_Y11F_ as 100% soluble underestimates degradation, rather than overestimating it.

We applied the correlation factor to 103 heterologous proteins to estimate how much expression was lost due to degradation for each protein. This analysis showed that, on average, 37% of protein expression was lost to degradation, and that 32 out of 103 proteins showing degradation of more than half of the produced protein. This is similar to a large-scale *E. coli* expression study, where approximately half of 7,733 proteins showed low to medium solubility^5^. Since only enzymes with previously demonstrated activity in cyanobacteria were included, proteins with very poor soluble expression were probably underrepresented in our dataset. We further investigated whether specific protein features could explain differences in soluble expression and found no correlation with eukaryotic or bacterial origin. We did, however, observe a weak correlation between protein size and soluble expression, consistent with previous studies showing that smaller proteins (∼20 kDa) generally have higher solubility than larger proteins (>40 kDa)^43^. Other studies have also identified amino acid composition as an important factor influencing soluble expression, where a higher proportion of hydrophilic and charged residues is associated with increased solubility, while stretches of hydrophobic amino acids negatively affect soluble expression^5,44^. However, we found no correlation between the GRAVY score, a measure of overall protein hydrophobicity, and soluble expression, suggesting that overall hydrophobicity alone does not explain the differences in expression observed in our dataset.

What is the effect of this protein wastage on the cell? Although degradation of recombinant proteins represents a waste of cellular resources, its direct metabolic cost is likely modest because amino acids can largely be recycled after proteolysis.

Nevertheless, synthesis of proteins that never reach a functional state still consumes ATP and occupies ribosomes that could otherwise be used for productive protein synthesis. The more important consequence for metabolic engineering, however, is the reduction in functional enzyme abundance. For enzymes that limit pathway flux, lower protein accumulation can directly reduce product formation, whereas improving protein stability may increase pathway flux without requiring stronger expression. This may be particularly relevant in cyanobacteria, where heterologous protein expression is generally lower than in *E. coli* and few promoters achieve very high expression levels^45^.

Reducing protein loss could therefore provide an alternative strategy for increasing enzyme abundance when further increases in expression are difficult to achieve. However, our results suggest that Clp repression is not necessarily an effective strategy for increasing enzyme activity, as increased protein abundance does not always translate into increased levels of functional enzyme.

We also found several examples of enzymes currently used in metabolic engineering applications that exhibit considerable expression limitations. For example, many enzymes from terpenoid biosynthetic pathways showed high levels of degradation. Many terpenoid substrates contain long hydrophobic carbon chains, and binding of these molecules often requires hydrophobic regions within the active site and substrate entry channels. This increased hydrophobicity may contribute to a higher tendency of these proteins to misfold or aggregate when expressed in a heterologous host.

Additionally, several enzymes identified as highly degraded in our dataset have previously shown signs of poor accumulation in cyanobacterial engineering studies. For example, aldehyde deformylating oxygenase (ADO) from *P. marinus* displayed a Clp ratio of 2.6, suggesting that more than 95% of the protein may be lost through degradation. Previous engineering studies in *Synechocystis* have reported very low ADO accumulation levels, but measurable product formation was still achieved, indicating that the remaining enzyme activity was sufficient to support the engineered pathway^46^. Another example is Dxs, the initial step of the MEP terpenoid pathway. Both Dxs variants tested in this study, from *E. coli* and *C. forskohlii*, showed low expression and substantial degradation. Previous studies have reported that *E. coli* Dxs is largely insoluble when overexpressed in its native host^47^, and similar limitations in soluble accumulation have been observed for native *Synechocystis* Dxs^48^. Dxs is also subject to feedback regulation by downstream metabolites, as IPP and DMAPP have been reported to promote Dxs aggregation^49^. Therefore, the low expression and high degradation observed for the Dxs variants may result from a combination of poor solubility of Dxs when overexpressed and aggregation triggered by feedback regulation.

If low soluble expression limits enzyme accumulation and pathway activity, one option to address this problem is to introduce solubility-enhancing mutations to the enzyme. Several computational tools have been developed to predict such variants, including PROSS^50^. However, improving protein stability does not necessarily improve enzyme performance. For example, a study in *Synechocystis* introduced PROSS-designed mutations in Pdc to improve ethanol production, but none of the engineered variants resulted in detectable ethanol production^51^. This highlights the risk that mutations intended to improve protein stability may also alter residues important for catalysis. In addition, our results demonstrate that replacing poorly accumulating enzymes with homologous variants can be an effective alternative, providing a simpler strategy to improve enzyme accumulation without requiring extensive protein redesign. This may represent the simplest approach when suitable homologs are available, although it is less applicable for enzymes with few naturally occurring alternatives.

In this study, we developed a method to assess protein degradation in *Synechocystis*. We found that a significant fraction of heterologously expressed proteins have unrealized expression, that is, a potential for higher expression if the proteins were more stable and were not targets of the protein quality control system. Addressing these limitations could improve enzyme expression and increase catalytic activity in metabolic pathways, leading to higher product titers.

## Materials and methods

### Cultivation conditions

*Synechocystis* sp. PCC 6803 was cultivated in 2x concentrated BG11 (Sigma-Aldrich), supplemented with 20 mM HEPES (Thermo Fisher) pH 7.5 and 3 g/L thiosulfate (Sigma-Aldrich). Cultures were grown at 30°C, 100 μmol photons/m^2^/s, atmospheric CO_2_ and 150 rpm in 4 mL cultures in 6-well tissue culture suspension plates (Sarstedt). Cultures were supplemented with 25 μg/mL kanamycin (Sigma-Aldrich) if they carried an expression plasmid and 12 μg/mL erythromycin (Thermo Scientific) if they had a CRISPRi knockdown cassette. For fluorescence measurements and protein sampling, fresh cultures were diluted to OD_730_ 0.1, grown for two days, induced with 1.25 μg/mL anhydrotetracycline (aTc, Sigma-Aldrich) or left uninduced, and grown for two additional days. On the fourth day, cultures were measured and sampled, unless otherwise stated.

### Cloning

Gene sequences were synthesized from IDT, ordered from Addgene or donated from other labs. Genes were cloned into plasmids pBG (Ptrc BCD) or pBG2 (Ptrc2O), both derived from pPMQAK1 (see Supplementary File 12-13 for plasmid sequences)^52^. *E. coli* DH5αZ1 (Expressys) was used for subcloning of plasmids and for *E. coli* expression measurements. Plasmids were introduced into *Synechocystis* by triparental mating as previously described^53^, using the pRL443 conjugative helper plasmid. Three colonies were picked per conjugated strain and colony PCR was used to verify the presence of the desired plasmid.

CRISPRi strains were constructed as described in Yao et al. (2015)^22^. A strain previously engineered with dCas9 integrated into *psbA1* was modified with two gRNA cassettes downstream of the dCas9 gene (Supplementary File 14). Sequence of gRNAs are available in Supplementary Table 2. The insertion of the gRNA cassettes was done using homologous recombination, using established protocols^54^. Colonies from transformations were screened by extracting genomic DNA using a GeneJet Genomic DNA Purification Kit (Thermo Scientific) and amplifying target region using PCR.

### Pigment analysis

Chlorophyll and carotenoids pigments were measured according to previous methods^55^. Phycobiliproteins were extracted and analyzed based on a modified version of a previously published method^56^. Culture samples corresponding to 2 mL of OD_730_ = 2.52 were pelleted by centrifugation at max speed (20,800 g) for 4 min at 4°C, and supernatant was removed. 100 μl of beads (Sigma-Aldrich G1277, 212-300 μm) were added together with 100 μL of ice-cold PBS. Cells were lysed using a FastPrep-24 5G bead beater (MP Biomedicals), six cycles of 45 s. 1 mL of PBS was added, and samples were vortexed and incubated on ice for 20 min. The extracted pigments were recovered by centrifuging the samples at max speed, 4°C for 5 min, and pigments were quantified using the absorbance of the supernatant at 615, 652 and 720 nm using a V-1200 spectrophotometer (VWR) with PBS used as blank. Phycocyanin content = ((A_615_ - A_720_) - 0.474 x (A_652_ - A_720_)) / 5.34 [mg/mL]. Allophycocyanin = ((A_652_ - A_720_) - 0.208 x (A_615_ - A_720_)) / 5.09 [mg/mL].

### Fluorescence measurements

Split-GFP fluorescence was measured by transferring 100 μl culture samples to 96-well microplates (Greiner Bio-One) and measuring fluorescence at 485 nm excitation and 535 nm emission using a SpectraMax i3x plate reader (Molecular Devices).

Fluorescence values were normalized to optical density measured at 730 nm for *Synechocystis* samples and 600 nm for *E. coli* samples. *Synechocystis* cultures were grown and sampled as described in the “Cultivation conditions” section. For *E. coli* measurements, frozen DH5αZ1 cultures containing the expression plasmids were streaked onto LB agar plates. Three colonies per strain were inoculated into 1 ml EZ-Rich medium (Teknova) in 12-well plates and grown overnight. The following day, cultures were reinoculated into fresh 1 ml EZ-Rich medium supplemented with 1 mM IPTG (Sigma-Aldrich) and grown at 37°C. After 24 h, 100 μl culture samples were transferred to 96-well microplates and fluorescence was measured as described above.

### Half-life measurements

Fresh cultures were inoculated to OD_730_ 0.1 in 6-well plates and grown for 24 hours. Next day, 200 μg/mL chloramphenicol (Tokyo chemical industry) and 200 μg/mL spectinomycin (Sigma-Aldrich) was added to the cultures. Fluorescence was measured every hour for seven hours. Half-life was calculated from fluorescence / OD_730_ values, by first dividing all values by the 0h value (N/N_0_), then taking the natural logarithm (ln(N/N_0_)) and from those values, calculating the slope when plotted against time (−k). Half-life was then calculated as t_1/2_ = ln(2)/k.

### Protein sampling and quantification

Proteins were extracted from 4 mL harvested cell pellets. Cells were lysed the same way as for pigment analysis, centrifuged at 20,800 g for 5 min, and the supernatant was collected. Total protein concentration in the cell-free extracts was determined using Bradford assay (Bio-Rad, cat#5000006). A standard curve was generated using known concentrations of bovine serum albumin (BSA) analyzed under identical assay conditions, and protein concentrations of the cell-free extracts were calculated from the slope of this standard curve.

### Proteomic sample preparation

Proteins were reduced in 10 mM DTT at 55°C for 45 min and alkylated in 17 mM IAA for 30 min in darkness. Proteins were digested overnight using Trypsin/Lys-C Protease Mix (Thermo Scientific) at a 1:50 protease:protein ratio. Digestion was quenched by addition of formic acid to pH ∼ 2 before desalting using stage tips packed with six layers of C18 Empore™ SPE Disks (Merck). Desalted peptides were dried in a Speed Vac at 45 °C and stored at −20 °C before LC-MS injection.

### Mass spectrometry method

Peptides were resuspended in 0.1% formic acid and analyzed using a Q-Exactive HF Hybrid Quadrupole-Orbitrap Mass Spectrometer operated in positive mode with an UltiMate 3000 RSLCnano System with an EASY-Spray ion source. Peptides were loaded on a C18 Acclaim PepMap 100 trap column (75 μm x 2 cm, 3 μm, 100 Å) at a flow rate of 7 μL per min solvent A (3% acetonitrile, 0.1% formic acid in water). An ES802 EASY-Spray PepMap RSLC C18 Column (75 μm x 25 cm, 2 μm, 100 Å) with a flow rate of 0.7 μL per minute was used to separate the peptides with a 40 minutes linear gradient from 1% to 32% solvent B (95% acetonitrile, 0.1% formic acid in water). Peptides were measured using one full scan (resolution 30,000 at 200 m/z, mass range 300–1200 m/z) followed by 30 MS2 DIA scans (resolution 30,000 at 200 m/z, mass range 350–1000 m/z) with an isolation window of 10 m/z. Precursor ion fragmentation was performed with high-energy collision-induced dissociation at an NCE of 26. Maximum injection times for the MS1 and MS2 were 105 ms and 55 ms, and automatic gain control was set to 3E6 and 1E6, respectively.

### Data analysis method

Proteomic data was searched using the EncyclopeDIA version 1.2.2 search engine^57^. The prosit intensity prediction model “Prosit_2020_intensity_hcd” was used to generate a predicted peptide library for *Synechocystis* from the UniProt reference proteome UP000001425. A maximum of two missed cleavages and a false discovery rate of 1% was used. Proteins were median-normalized and fold-changes were calculated using the MSstats (v.4.4.1) package in R^58^. The p-values were adjusted for multiple hypothesis testing using the Benjamini-Hochberg method, with an adjusted p-value threshold of 0.05. Proteins detected in fewer than 2 replicates were excluded from statistical analysis.

### Western blot

An equal amount of cells was taken from each culture, corresponding to 2 mL of OD_730_ of 1. Cells were lysed as described in the “Protein extraction” section, but samples were not centrifuged after, to prevent removal of aggregated proteins. Samples representing total protein fraction were run on two Mini-PROTEAN TGX Stain-Free gels (BIO-RAD), one for western blotting and one stained with QC Colloidal Coomassie Stain (BIO-RAD) as equal loading control. Western blot analysis was performed as described in Hoffman et al. (2025)^59^. N-terminal strep-tag of AcP, Im7 and Tim was detected using an antistrep-tag II rabbit IgG antibody (ab76949) together with the secondary horseradish peroxidase conjugated goat antirabbit IgG antibody (Abcam #ab672).

### Enzymatic Activity Assay

Enzymatic activity of the cell-free extracts was assessed spectrophotometrically by monitoring NADPH (Roche, 2646-71-1) consumption at 340 nm using a plate reader, SpectraMax i3X. For each assay, a cell-free extract corresponding to 100 µg total protein was added to a 200 µL reaction mixture made of 50 mM phosphate buffer (pH 6.4), 0.8 mM NADPH, and 1 mM cyclohexanone. Prior to the addition of NADPH and cyclohexanone, the reaction buffer was continuously purged with N₂ gas to maintain anaerobic conditions. Reactions were carried out in 96-well plates and monitored continuously for 12 min. Activity was determined for all prepared cell-free extracts, with each strain assayed in biological triplicate. Reaction rates were calculated from the slope of the absorbance decrease at 340 nm for each sample and normalized to protein concentration and reaction time to obtain reaction rates. A standard curve was generated by serial dilution of NADPH (0–1 mM) to correlate absorbance at 340 nm with NADPH concentration. This standard curve was used to convert measured reaction rates into NADPH consumption rates.

## Acknowledgement

This work was supported by Formas Mobility Grant Nr. 2017–00335, Novo Nordisk Foundation Postdoctoral Fellowship Nr. NNF22C0079474 and Swedish Foundation for Strategic Research SSF, grant no. ARC19-0051. The project is co-funded by the European Union under project Solar to Butanol S2B, Grant agreement ID: 101172911. We also would like to thank the lab of Professor Robert Kourist and Professor Pia Lindberg for donating plasmids.

## Author contributions

E.E. conceived and designed the study, performed the majority of the experiments, analyzed and interpreted the data, and wrote the manuscript. H.M. performed the *E. coli* expression measurements and *in vitro* enzymatic assays and reviewed and commented on the manuscript. A.K. performed the proteomic experiments and proteomic data analysis and reviewed and commented on the manuscript. A.D. contributed to the *E. coli* expression measurements. E.P.H. contributed to supervision and scientific discussion of the study and reviewed and commented on the manuscript. All authors approved the final manuscript.

## References

1. Singh, V. et al. Strategies for Fermentation Medium Optimization: An In-Depth Review. Front. Microbiol. 7, (2017).

2. Jaiswal, D., Sahasrabuddhe, D. & Wangikar, P. P. Cyanobacteria as cell factories: the roles of host and pathway engineering and translational research. Current Opinion in Biotechnology 73, 314–322 (2022).

3. Liu, X., Tang, K. & Hu, J. Application of Cyanobacteria as Chassis Cells in Synthetic Biology. Microorganisms 12, 1375 (2024).

4. Angermayr, S. A. et al. Exploring metabolic engineering design principles for the photosynthetic production of lactic acid by Synechocystis sp. PCC6803. Biotechnol Biofuels 7, 99 (2014).

5. Price, W. N. et al. Large-scale experimental studies show unexpected amino acid effects on protein expression and solubility in vivo in E. coli. Microb Informatics Exp 1, 6 (2011).

6. Luan, C.-H. et al. High-Throughput Expression of C. elegans Proteins. Genome Res. 14, 2102–2110 (2004).

7. DePristo, M. A., Weinreich, D. M. & Hartl, D. L. Missense meanderings in sequence space: a biophysical view of protein evolution. Nat Rev Genet 6, 678–687 (2005).

8. Györkei, Á. et al. Proteome-wide landscape of solubility limits in a bacterial cell. Sci Rep 12, 6547 (2022).

9. Mogk, A., Bukau, B. & Kampinga, H. H. Cellular Handling of Protein Aggregates by Disaggregation Machines. Molecular Cell 69, 214–226 (2018).

10. Zhang, X., Betterle, N., Hidalgo Martinez, D. & Melis, A. Recombinant Protein Stability in Cyanobacteria. ACS Synth. Biol. 10, 810–825 (2021).

11. Bouchnak, I. & Van Wijk, K. J. Structure, function, and substrates of Clp AAA+ protease systems in cyanobacteria, plastids, and apicoplasts: A comparative analysis. Journal of Biological Chemistry 296, 100338 (2021).

12. Imai, K., Kitayama, Y. & Kondo, T. Elucidation of the Role of Clp Protease Components in Circadian Rhythm by Genetic Deletion and Overexpression in Cyanobacteria. Journal of Bacteriology 195, 4517–4526 (2013).

13. Cheregi, O., Wagner, R. & Funk, C. Insights into the Cyanobacterial Deg/HtrA Proteases. Front. Plant Sci. 7, (2016).

14. Kamata, T. et al. Quality control of Photosystem II: an FtsH protease plays an essential role in the turnover of the reaction center D1 protein in Synechocystis PCC 6803 under heat stress as well as light stress conditions. Photochem Photobiol Sci 4, 983–990 (2005).

15. Foit, L. et al. Optimizing Protein Stability In Vivo. Molecular Cell 36, 861–871 (2009).

16. Ren, C., Wen, X., Mencius, J. & Quan, S. An enzyme-based biosensor for monitoring and engineering protein stability in vivo. Proceedings of the National Academy of Sciences 118, e2101618118 (2021).

17. Sullivan, B. J. et al. Stabilizing Proteins from Sequence Statistics: The Interplay of Conservation and Correlation in Triosephosphate Isomerase Stability. Journal of Molecular Biology 420, 384–399 (2012).

18. Cabantous, S., Terwilliger, T. C. & Waldo, G. S. Protein tagging and detection with engineered self-assembling fragments of green fluorescent protein. Nat Biotechnol 23, 102–107 (2005).

19. Kamiyama, D. et al. Versatile protein tagging in cells with split fluorescent protein. Nat Commun 7, 11046 (2016).

20. Chiti, F. et al. Mutational analysis of acylphosphatase suggests the importance of topology and contact order in protein folding. Nat Struct Mol Biol 6, 1005–1009 (1999).

21. Capaldi, A. P., Kleanthous, C. & Radford, S. E. Im7 folding mechanism: misfolding on a path to the native state. Nat Struct Mol Biol 9, 209–216 (2002).

22. Yao, L., Cengic, I., Anfelt, J. & Hudson, E. P. Multiple Gene Repression in Cyanobacteria Using CRISPRi. ACS Synth. Biol. 5, 207–212 (2016).

23. Huang, H.-H. & Lindblad, P. Wide-dynamic-range promoters engineered for cyanobacteria. J Biol Eng 7, 10 (2013).

24. Barker, M., de Vries, R., Nield, J., Komenda, J. & Nixon, P. J. The Deg Proteases Protect *Synechocystis* sp. PCC 6803 during Heat and Light Stresses but Are Not Essential for Removal of Damaged D1 Protein during the Photosystem Two Repair Cycle*. Journal of Biological Chemistry 281, 30347–30355 (2006).

25. Giner-Lamia, J. et al. Identification of the direct regulon of NtcA during early acclimation to nitrogen starvation in the cyanobacterium Synechocystis sp. PCC 6803. Nucleic Acids Res 45, 11800–11820 (2017).

26. Krauspe, V., Timm, S., Hagemann, M. & Hess, W. R. Phycobilisome Breakdown Effector NblD Is Required To Maintain Cellular Amino Acid Composition during Nitrogen Starvation. Journal of Bacteriology 204, e00158–21 (2022).

27. Liu, X. et al. Interactome Analysis of ClpX Reveals Its Regulatory Role in Metabolism and Photosynthesis in Cyanobacteria. J. Proteome Res. 23, 1174–1187 (2024).

28. Liu, X., Miao, R., Lindberg, P. & Lindblad, P. Modular engineering for efficient photosynthetic biosynthesis of 1-butanol from CO_2_ in cyanobacteria. Energy & Environmental Science 12, 2765–2777 (2019).

29. Englund, E., Shabestary, K., Hudson, E. P. & Lindberg, P. Systematic overexpression study to find target enzymes enhancing production of terpenes in Synechocystis PCC 6803, using isoprene as a model compound. Metabolic Engineering 49, 164–177 (2018).

30. Martin, V. J. J., Pitera, D. J., Withers, S. T., Newman, J. D. & Keasling, J. D. Engineering a mevalonate pathway in Escherichia coli for production of terpenoids. Nat Biotechnol 21, 796–802 (2003).

31. Mutalik, V. K. et al. Precise and reliable gene expression via standard transcription and translation initiation elements. Nat Methods 10, 354–360 (2013).

32. Lou, C., Stanton, B., Chen, Y.-J., Munsky, B. & Voigt, C. A. Ribozyme-based insulator parts buffer synthetic circuits from genetic context. Nat Biotechnol 30, 1137–1142 (2012).

33. Wang, J. et al. Research on enhancing the expression and immobilization of oxalate decarboxylase via the bicistronic translation coupling strategy. Journal of Cleaner Production 527, 146650 (2025).

34. Ranaghan, M. J., Li, J. J., Laprise, D. M. & Garvie, C. W. Assessing optimal: inequalities in codon optimization algorithms. BMC Biol 19, 36 (2021).

35. Gupta, M. et al. Global protein turnover quantification in Escherichia coli reveals cytoplasmic recycling under nitrogen limitation. Nat Commun 15, 5890 (2024).

36. Varshavsky, A. The N-end rule: functions, mysteries, uses. Proc. Natl. Acad. Sci. U.S.A. 93, 12142–12149 (1996).

37. Ratelade, J. et al. Production of Recombinant Proteins in the lon-Deficient BL21(DE3) Strain of Escherichia coli in the Absence of the DnaK Chaperone. Applied and Environmental Microbiology 75, 3803–3807 (2009).

38. Shestakov, S. V. & Khyen, N. T. Evidence for genetic transformation in blue-green alga Anacystis nidulans. Molec. Gen. Genet. 107, 372–375 (1970).

39. Mutalik, V. K. et al. Quantitative estimation of activity and quality for collections of functional genetic elements. Nat Methods 10, 347–353 (2013).

40. Pohl, S. et al. Proteomic analysis of Bacillus subtilis strains engineered for improved production of heterologous proteins. PROTEOMICS 13, 3298–3308 (2013).

41. Liu, X., Meng, L., Wang, X., Yang, Y. & Bai, Z. Effect of Clp protease from *Corynebacterium glutamicum* on heterologous protein expression. Protein Expression and Purification 189, 105928 (2022).

42. Francis, D. M. & Page, R. Strategies to Optimize Protein Expression in E. coli. Current Protocols in Protein Science 61, 5.24.1–5.24.29 (2010).

43. Dyson, M. R., Shadbolt, S. P., Vincent, K. J., Perera, R. L. & McCafferty, J. Production of soluble mammalian proteins in Escherichia coli: identification of protein features that correlate with successful expression. BMC Biotechnol 4, 32 (2004).

44. Goh, C.-S. et al. Mining the Structural Genomics Pipeline: Identification of Protein Properties that Affect High-throughput Experimental Analysis. Journal of Molecular Biology 336, 115–130 (2004).

45. Zhou, J. et al. Discovery of a super-strong promoter enables efficient production of heterologous proteins in cyanobacteria. Sci Rep 4, 4500 (2014).

46. Yunus, I. S. et al. Synthetic metabolic pathways for conversion of CO2 into secreted short-to medium-chain hydrocarbons using cyanobacteria. Metabolic Engineering 72, 14–23 (2022).

47. Zhou, K., Zou, R., Stephanopoulos, G. & Too, H.-P. Enhancing solubility of deoxyxylulose phosphate pathway enzymes for microbial isoprenoid production. Microb Cell Fact 11, 148 (2012).

48. Kudoh, K. et al. Overexpression of endogenous 1-deoxy-d-xylulose 5-phosphate synthase (DXS) in cyanobacterium *Synechocystis* sp. PCC6803 accelerates protein aggregation. Journal of Bioscience and Bioengineering 123, 590–596 (2017).

49. Di, X., et al. MEP pathway products allosterically promote monomerization of deoxy-D-xylulose-5-phosphate synthase to feedback-regulate their supply. Plant Comm 4, (2023).

50. Goldenzweig, A. et al. Automated Structure- and Sequence-Based Design of Proteins for High Bacterial Expression and Stability. Molecular Cell 63, 337–346 (2016).

51. Nishiguchi, H., Niide, T., Toya, Y. & Shimizu, H. Efficient selection of pyruvate decarboxylase sequences from database for high ethanol productivity in *Synechocystis* sp. PCC 6803. Journal of Bioscience and Bioengineering 140, 123–131 (2025).

52. Huang, H.-H., Camsund, D., Lindblad, P. & Heidorn, T. Design and characterization of molecular tools for a Synthetic Biology approach towards developing cyanobacterial biotechnology. Nucleic Acids Res 38, 2577–2593 (2010).

53. Elhai, J. & Wolk, C. P. Conjugal Transfer of DNA to Cyanobacteria In Methods Enzymol. (Academic Press, 1988).

54. Heidorn, T. et al. Synthetic biology in cyanobacteria: engineering and analyzing novel functions. in Methods in enzymology vol. 497 539–579 (Elsevier, 2011).

55. Zavřel, T., Sinetova, M. A. & Červený, J. Measurement of Chlorophyll a and Carotenoids Concentration in Cyanobacteria. Bio-protocol 5, (2015).

56. Zavřel, T., Chmelík, D., Sinetova, M. A. & Červený, J. Spectrophotometric Determination of Phycobiliprotein Content in Cyanobacterium Synechocystis. Journal of Visualized Experiments (JoVE) e58076 (2018) doi:10.3791/58076.

57. Searle, B. C. et al. Chromatogram libraries improve peptide detection and quantification by data independent acquisition mass spectrometry. Nat Commun 9, 5128 (2018).

58. Choi, M. et al. MSstats: an R package for statistical analysis of quantitative mass spectrometry-based proteomic experiments. Bioinformatics 30, 2524–2526 (2014).

59. Hoffmann, U. A. et al. A Cyanobacterial Screening Platform for Rubisco Mutant Variants. ACS Synth. Biol. 14, 2619–2633 (2025).

