## Supplementary Information for "Expression landscape of metabolic engineering enzymes in cyanobacteria"

for

Supplementary Figure 1 - Growth curves of ClpS1/S2 and Deg/HtrA CRISPRi knockdown.

Supplementary Figure 2 - Volcano plots of ClpS1/S2 and Deg/HtrA knockdown.

Supplementary Figure 3 - Pigment analysis of Clp knockdown strains.

Supplementary Figure 4 - Expression of AcP variants in ClpS1/S2 and Deg/HtrA knockdown.

Supplementary Table 1 - Comparison of ClpX interaction partners with proteins affected by Clp knockdown.

Supplementary Table 2 - Guide RNA sequences for dCas9 knockdown.

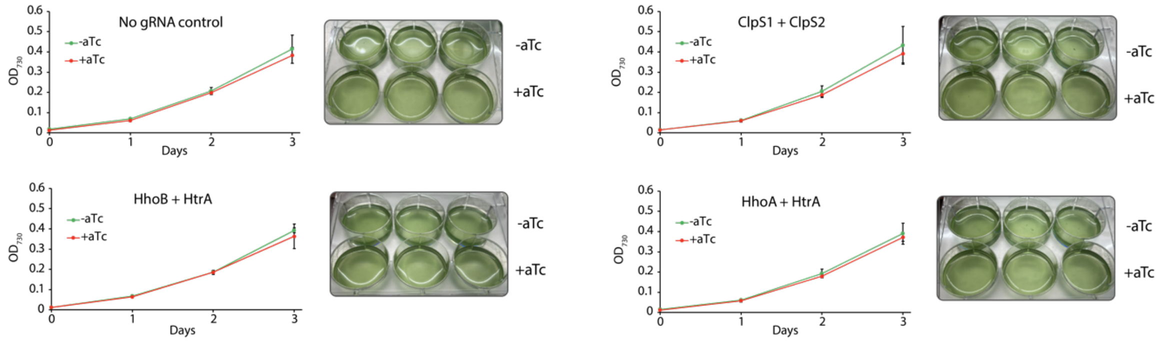


**Supplementary Figure 1**. Growth curves of ClpS1/S2 and Deg/HtrA CRISPRi knockdown. Inserted pictures show the colour of the culture. Optical density was measured using a plate reader which gives $OD_{730}$ values ~5 times lower than when measured with a standard 1 cm pathlength cuvette spectrophotometer. Error bars represents the standard deviation of three biological replication.

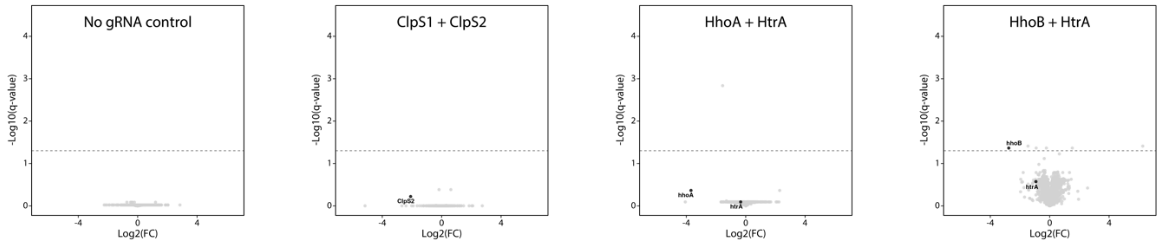


**Supplementary Figure 2**. Volcano plots of ClpS1/S2 and Deg/HtrA knockdown. Targets of knockdown are color coded black, phycobiliproteins are blue, proteases are red and chaperones/heat shock proteins are green. Results are based on three biological replicates.

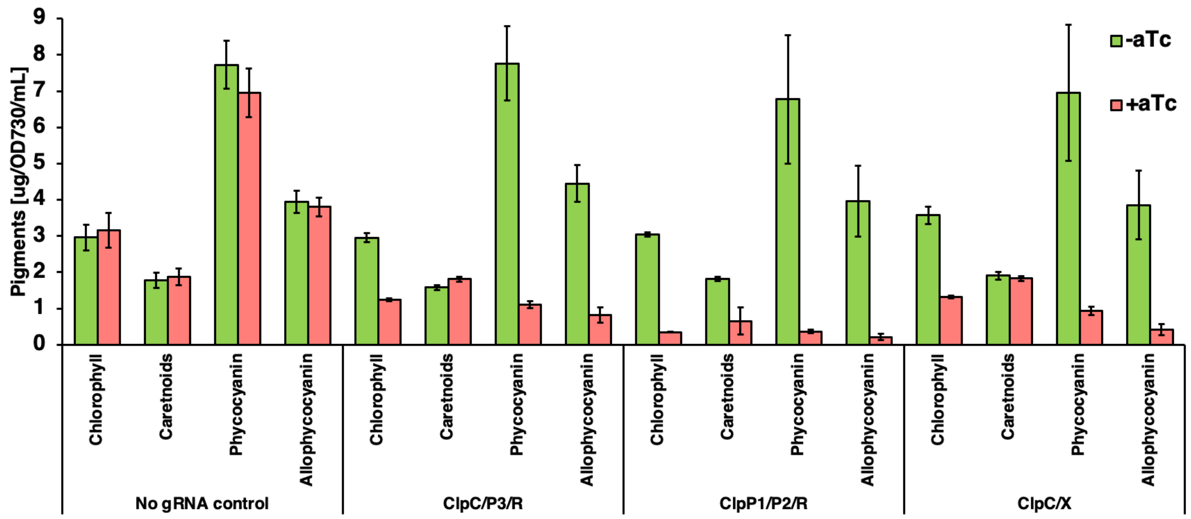


**Supplementary Figure 3**. Pigment analysis of Clp knockdown strains. Results are based on three biological replicates, error bars represent standard deviation.

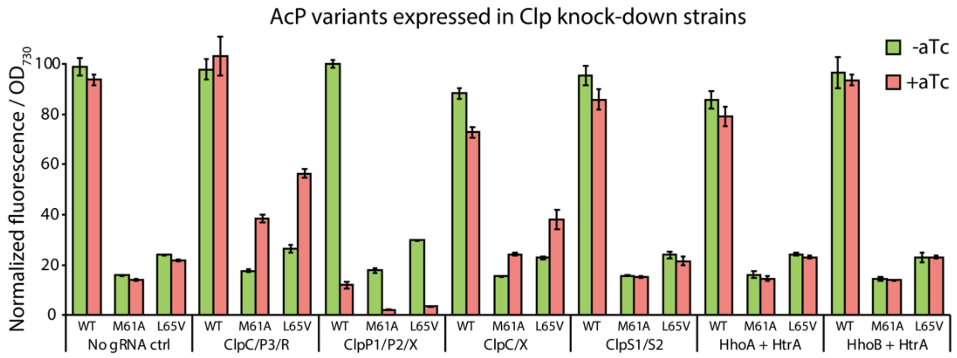


**Supplementary Figure 4**. Expression of AcP variants in Clp knockdown strains. Error bars represents the standard deviation of three biological replication.

**Supplementary Table 1**. Comparison of ClpX interaction partners identified in Liu et al. 2024 with proteins affected by Clp knockdown in this study. Significance was determined based on a $\log_2$ fold change >1 and adjusted p-value <0.05.

| KD strain | Protein set | Total Proteins | Not significantly changed | Significantly changed | Increased abundance | Decreased abundance |
|---|---|---|---|---|---|---|
| ClpC/P3/R | Previously identified ClpX interactors | 503 | | | | |
| | All proteins detected in KD strain | 2521 | 1724 | 797 | 379 | 418 |
| | Previously identified ClpX interactors detected | 430 | 316 | 114 | 39 | 75 |

| KD strain | Protein set | Total Proteins | Not significantly changed | Significantly changed | Increased abundance | Decreased abundance |
|---|---|---|---|---|---|---|
| ClpP1/P2/X | Previously identified ClpX interactors | 503 | | | | |
| | All proteins detected in KD strain | 2366 | 1740 | 626 | 436 | 190 |
| | Previously identified ClpX interactors detected | 424 | 286 | 138 | 9 | 129 |

| KD strain | Protein set | Total Proteins | Not significantly changed | Significantly changed | Increased abundance | Decreased abundance |
|---|---|---|---|---|---|---|
| ClpC/X | Previously identified ClpX interactors | 503 | | | | |
| | All proteins detected in KD strain | 2465 | 1716 | 749 | 345 | 404 |
| | Previously identified ClpX interactors detected | 424 | 291 | 133 | 27 | 106 |

**Supplementary Table 2**. Sequences of guide RNAs used together with dCas9. Each strain expressed two separate gRNAs.

| *Strain* | *Target 1* | *gRNA 1* | *Target 2* | *gRNA 2* |
|---|---|---|---|---|
| *ClpC/P3/R* | ClpC | gagcaagcatgattactttaa | ClpR | atgctgtcaggttggggtcga |
| *ClpP1/P2/X* | ClpP2 | aacgggagttgaccatgggtt | ClpP1 | acggttggaatcatggggaga |
| *ClpC/X* | ClpC | gagcaagcatgattactttaa | ClpX | attaatttacggacttgctcc |
| *ClpP1/P3/R* | ClpR | atgctgtcaggttggggtcga | ClpP1 | acggttggaatcatggggaga |
| *ClpS1/S2* | ClpS1 | ttgctaatcacgacagtaatc | ClpS2 | tgctgggcttgtttaaaactt |
| *HhoA/HtrA* | HhoA | gtctccaaatacgcattagca | HtrA | ggagcaattgggaaaacggct |
| *HhoB/HtrA* | HhoB | atgccttcagatggattgcca | HtrA | ggagcaattgggaaaacggct |
